# The effects of Alcohol Dependence on the CSF Proteome in Mice: Evidence for Blood-Brain Barrier Dysfunction and Neuroinflammation

**DOI:** 10.1101/2025.11.10.687295

**Authors:** Natalie P. Turner, Michal Bajo, Amanda J Roberts, Marisa Roberto, John R. Yates

## Abstract

Alcohol use disorder (AUD) represents a significant neurological health burden, yet the biological mechanisms underlying alcohol-induced brain pathology remain incompletely understood. Moreover, the molecular underpinnings of the transition from alcohol exposure to alcohol dependence are not well-characterized. We used mass spectrometry (MS)-based proteomics in a preliminary discovery study to compare cerebrospinal fluid (CSF) of alcohol-exposed Non-dependent (Non-dep) versus alcohol-dependent (Dep) mice that underwent the chronic intermittent ethanol (alcohol) – two-bottle choice (CIE-2BC) procedure and systemic anti-IL-6 Receptor antibody administration. CSF samples from individual mice were processed for proteomic analysis and digested with trypsin overnight. Peptides were analyzed via data-independent acquisition (DIA)-MS and data were processed in DIA-NN at 1% FDR. We identified 611 unique proteins across both groups, with 140 proteins differentially detected in CSF from Dep mice and 67 proteins specific to alcohol-exposed but Non-dep controls. The Dep-specific proteins revealed signatures of blood-brain barrier (BBB) disruption, neuroinflammation, cellular stress responses, and complement system activation. In contrast, Non-dep-specific proteins indicated preserved protective mechanisms including complement regulation, anti-inflammatory signaling, and neuronal calcium homeostasis. Ethanol-dependent-specific findings include MMP2, BIP, and to a lesser extent VE-cadherin (CDH5) and VCAM1, indicative of the beginnings of endothelial damage and BBB disruption, alongside established neuroinflammation markers GFAP, CHI3L1, and CX3CL1. This work provides novel preliminary protein-level evidence that alcohol exposure and alcohol dependence are dichotomous; despite the small sample size and limited power for moderate effect sizes, there appears to be a clear molecular transition from maintained protective mechanisms to vascular damage, BBB breakdown, and sustained neuroinflammation.

## Introduction

Alcohol use disorder (AUD) leads to significant neurological impairment, including cognitive decline, increased neuroinflammation, and blood-brain barrier (BBB) dysfunction. Chronic alcohol exposure disrupts tight junction proteins through multiple pathways, including matrix metalloproteinase activation, oxidative stress, and novel purinergic signaling mechanisms ^1,2^. These processes create a ‘leaky’ barrier that allows peripheral inflammatory mediators to enter the brain, potentially establishing the neuroinflammatory cycles observed in alcohol dependence, which are comparable to that seen in other neurological conditions such as Alzheimer’s Disease ^3,4^. There is accumulating evidence that cytokines can also be produced locally in the brain, offering an alternative hypothesis of compartmentalized neuroinflammation ^5^. Recent neuroimaging and cerebrospinal fluid studies have confirmed that AUD can be characterized by activated microglia, elevated cytokines, and complement system dysregulation ^6,7^. However, the molecular mechanisms distinguishing chronic alcohol exposure with moderate alcohol drinking from chronic alcohol exposure with excessive alcohol drinking that characterized alcohol dependence remain poorly understood. Current biomarker approaches focus primarily on hepatic dysfunction and chronic consumption detection ^8^, leaving a significant gap in neurological diagnostic and prognostic tools. In this study, we used the chronic intermittent ethanol vapor – two-bottle choice (CIE-2BC) procedure ^9–14^ to induce ethanol/alcohol dependence. Compared to moderate ethanol-drinking (2BC) mice that did not undergo ethanol vapor (defined as Non-dep) group, mice exposed to CIE-2BC escalated ethanol drinking, an indication of ethanol dependence (defined as Dep group).

Proteomic analysis of cerebrospinal fluid (CSF) offers a unique window into brain pathology by allowing detection of proteins normally confined to specific brain regions. Advances in biomarker research have established several proteins as reliable indicators of neurological damage, including glial fibrillary acidic protein (GFAP) for astrocyte activation ^15^ and ubiquitin C-terminal hydrolase L1 (UCHL1; gene = *Park5*) for neuronal injury ^16^. Despite these advances, a fundamental question remains unanswered: does alcohol dependence represent a quantitative progression from exposure, or does it involve qualitatively distinct pathophysiological mechanisms? This distinction has critical implications for early intervention strategies and biomarker development.

To address this question, we systematically evaluated and compared CSF samples collected from Non-dep and Dep mice by mass spectrometry (MS)-based proteomics. The CSF proteome of 2BC Non-dep mice was distinct from Dep mice, which contained proteins associated with neuroinflammation, compromised immune surveillance and loss of protective function. Additionally, the identification of the clinically validated biomarker of astrocyte activation, GFAP, and the protein Matrix Metalloproteinase 2 (MMP2) associated with BBB breakdown, in Dep mice supports the hypothesis that BBB disruption and immune dysregulation is unique to alcohol dependence.

## Results and Discussion

### Proteomic Profiles Differ Between Alcohol-Dependent and Non-Dependent Mice

Data-independent acquisition (DIA)-MS enables proteomic analysis of complex samples through simultaneous fragmentation of all peptides within a specific mass-to-charge ratio (*m*/*z*) isolation ‘window’, and sequential “stepping” through the entire MS1 mass range (e.g., 400 – 1200 *m*/*z*) with wide sequential windows (5 – 25 *m*/*z)*^17^. Because DIA is used to fragment all peptides within each isolation window rather than preferentially selecting the top N most abundant peptide precursors for fragmentation (as in data-dependent acquisition; DDA), this approach was selected for the current study to mitigate the possibility of missing peptide features. If features are missing in a DIA experiment, it is likely due to extremely low peptide abundance; thus, DIA provides a more confident analysis of qualitative datasets than in DDA experiments, where peptide features can be missing at random. Proteomic analysis using DIA-MS identified an average of 1816 peptide precursors and 330 proteins in the alcohol-exposed control group at 1% FDR (Figure 1a). In the ethanol dependent group, an average of 2667 peptide precursors and 466 proteins were identified. Further analysis found 140 proteins preferentially detected in alcohol-dependent mice and 67 proteins preferentially detected in Non-dep mice (Figure 1b). Given the limited statistical power for detecting moderate effect sizes (38% power for 75% vs 20% detection rates), the 140 Dep-specific and 67 Non-dep-specific proteins likely represent minimum estimates of the true proteomic differences between groups. A principal component analysis (PCA) plot with 95% confidence ellipse was generated to assess the relationships between replicates within and between groups (Figure 1c). PCA preserves the overall data structure while performing dimensionality reduction, enabling visualization of the relationships between data points (replicates) in two dimensions. The Dep animals formed a distinct cluster regardless of sex (orange points), whereas the Non-dependent animals were dispersed across the plot but could be distinguished along the PC2 axis by sex. One Non-dep animal clustered within the Dep group, which could indicate a potential outlier in the Non-dep group. The protein profiles of both groups were interrogated further using enrichment analysis and STRING network analysis to identify putative molecular mechanisms associated with alcohol-dependence.

**Figure 1:**
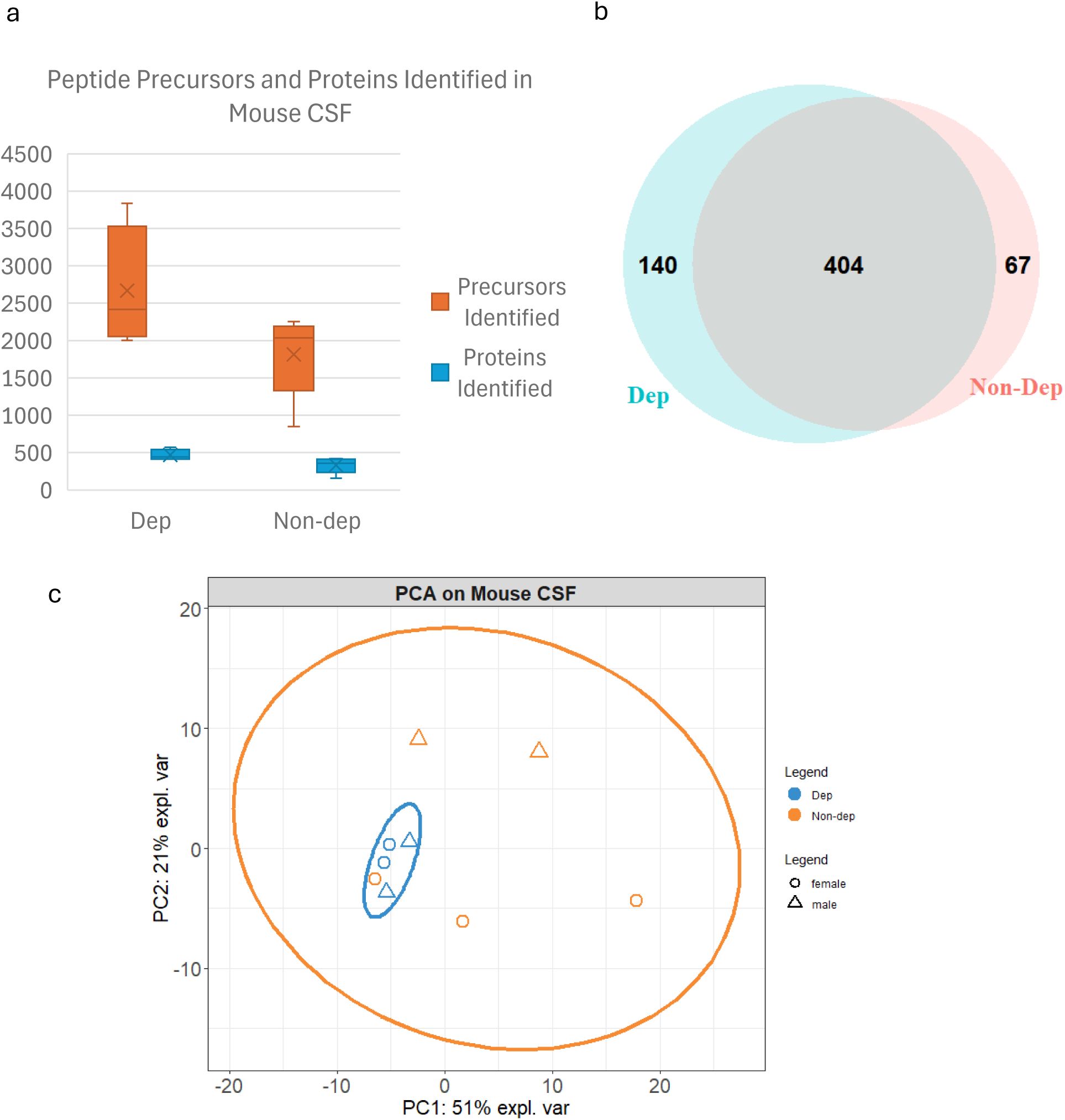
Proteomic analysis of Mouse CSF. a) Peptide precursors (orange) and proteins (blue) identified in Non-dep (*n* = 5) and Dep groups (*n* = 4). Box and whiskers plots; edges show the 25^th^ and 75^th^ percentile range; X represents the mean value, horizontal lines across boxes represents the median value, and the error bars indicate the minimum and maximum values. b) Venn diagram of proteins identified in Non-dep and Dep groups; Dep unique = 140; Non-dep unique = 67; overlap = 404. c) Principal component analysis (PCA) plot with 95% confidence ellipse of Non-dep (orange) and Dep (blue) mice. Dep mice show clustering regardless of sex, whereas control mice are more dispersed, although segregate by sex along the PC2 axis. Circles = females; triangles = males.

The top 10 group-specific proteins are shown in Figure 2. The heatmap displays segregation of groups based on treatment-type, demonstrating how a shift to alcohol dependency is reflected in the CSF proteome. Numerous proteins detected preferentially in the Dep group are associated with neuroinflammation (GFAP), and with synaptic and extracellular matrix (ECM) remodeling (Reelin – RELN; MMP2). The preferential detection of proteins such as CALB1 (Calbindin-1), SUMO2/SUMO3, and TAGL3 (Transgelin-3) suggests functional homeostatic and neuroprotective mechanisms are maintained in the Non-dep group. Additionally, detection patterns of proteins in immunoglobulin variable regions (KV5AB, KV5A7, LV1C, HVM02/03) indicate that immune surveillance may be lost in the Dep group. (For the full list of filtered protein identifications, see Supplementary file 1.)

**Figure 2:**
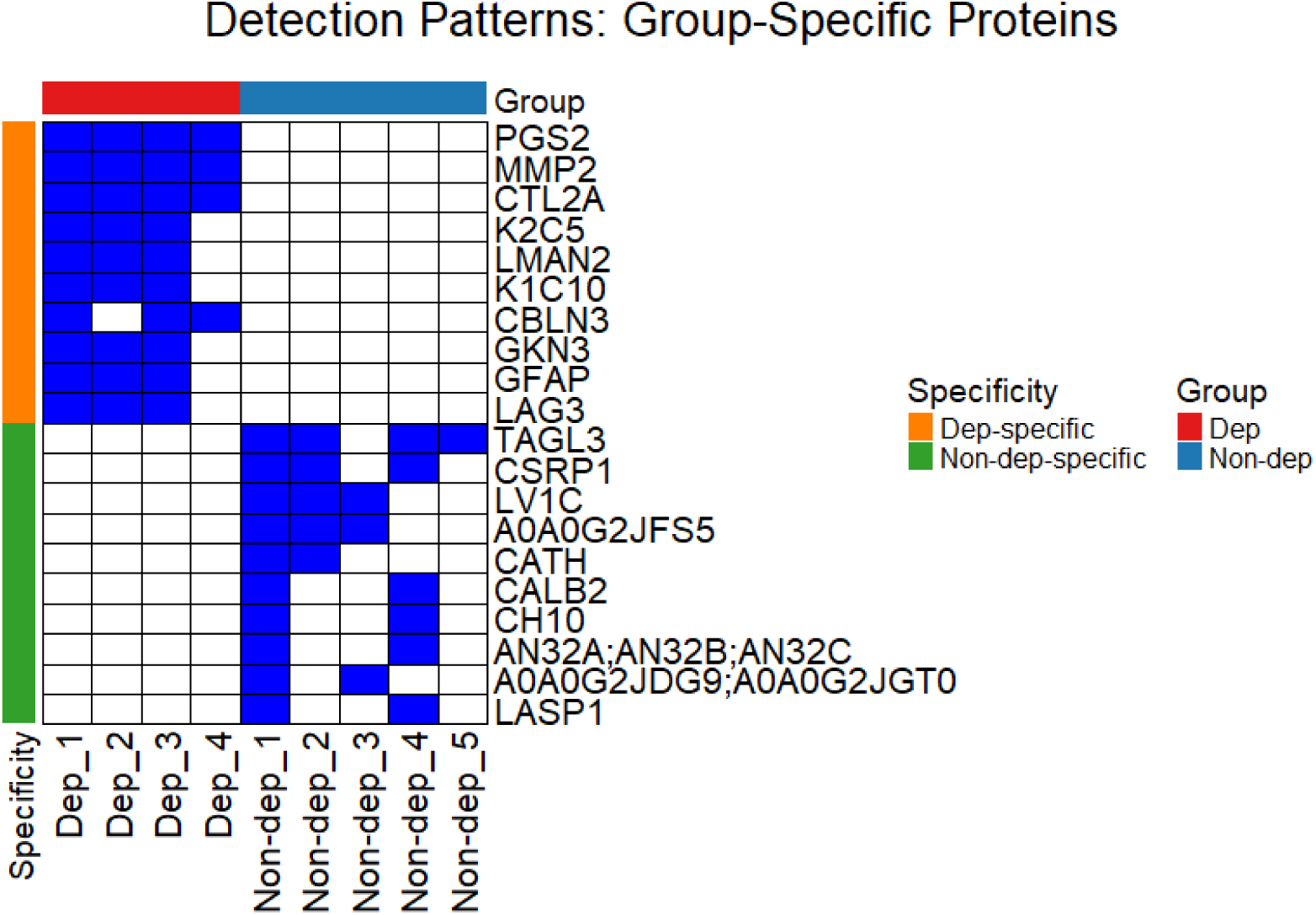
Detection Patterns of Group-Specific Proteins in Mouse CSF Samples. Heatmap showing the top 10 group-specific proteins detected (blue) or not-detected (white) across experimental samples. Samples were analyzed by data-independent acquisition (DIA) MS, therefore missing values are likely to represent protein levels below the lower limit of detection (i.e., not missing at random). Group specificity for the top 10 proteins was determined by protein detection in ≥ 2 of the replicates within each group and 0 replicates of the opposite group.

### Classification of Proteins Associated with Alcohol Dependence

The following criteria were used to classify protein-level evidence of alcohol dependence or non-dependence, based on detection patterns in mouse CSF samples:

**Strong Evidence:**

- Dep-preferred: ≥3/4 (≥75%) Dep AND ≤1/5 (≤20%) Non-dep
- Non-dep-preferred: ≥4/5 (≥80%) Non-Dep AND ≤1/4 (≤25%) Dep

**Moderate – Weak Evidence:**

- Dep-preferred: ≥2/4 (≥50%) Dep with clear preference
- Non-dep-preferred: ≥3/5 (≥60%) Non-dep with clear preference
- Weak: Detected in <50% of preferred group or substantial detection in both groups

Dep-preferred and Non-dep-preferred proteins with strong-moderate evidence are listed in Tables 1 and 2.

**Table 1:**
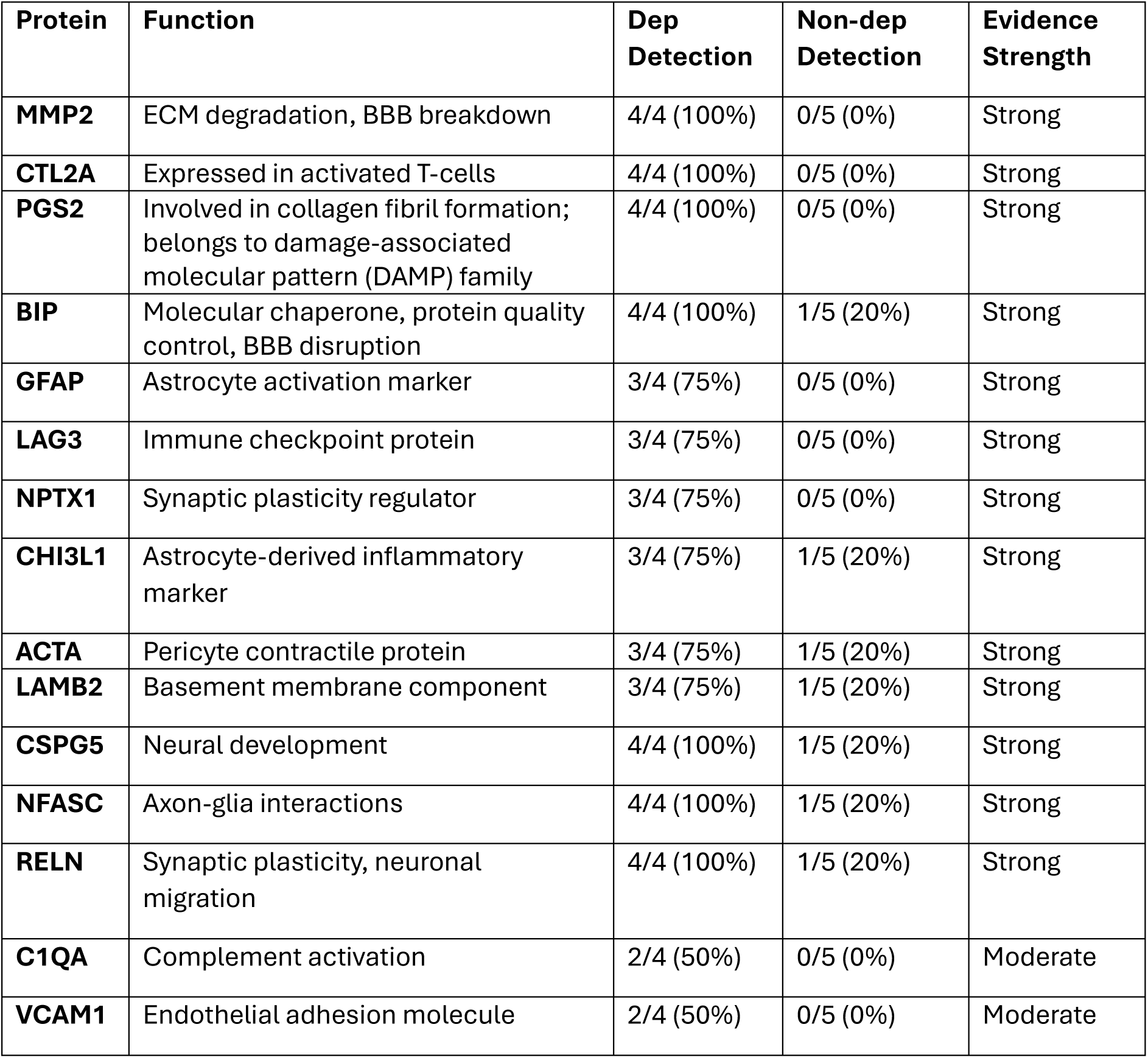
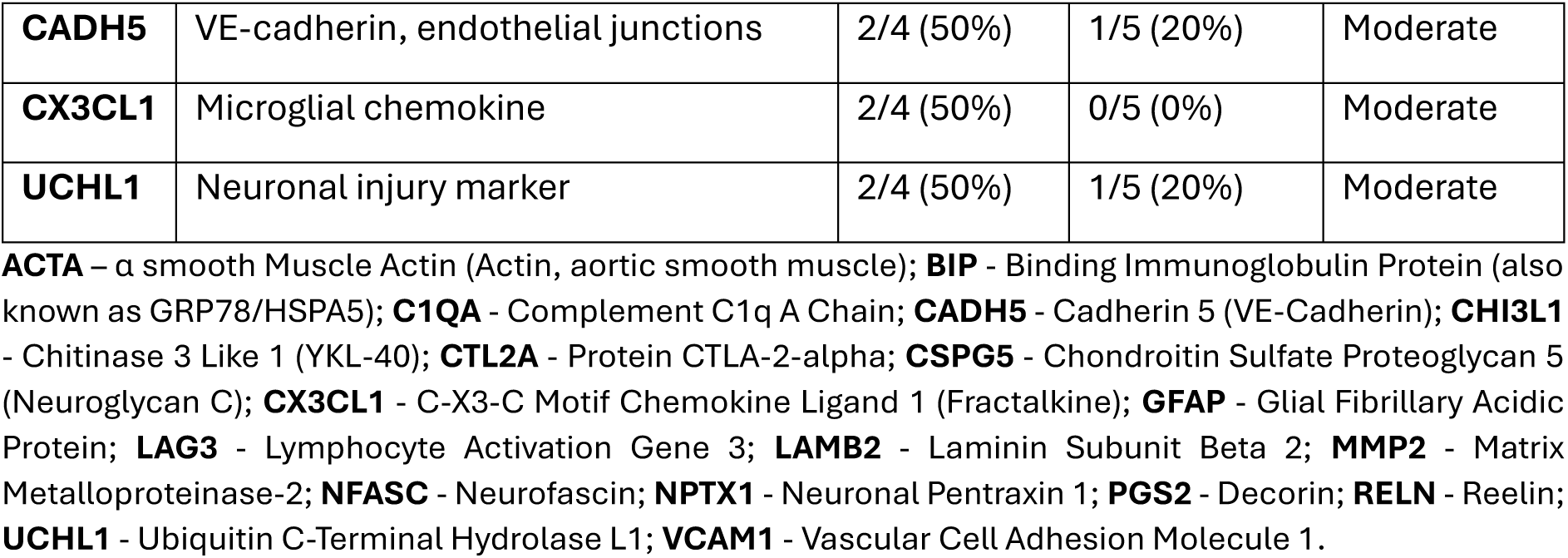
Dep-preferred proteins.

| Protein | Function | Dep Detection | Non-dep Detection | Evidence Strength |
| --- | --- | --- | --- | --- |
| **MMP2** | ECM degradation, BBB breakdown | 4/4 (100%) | 0/5 (0%) | Strong |
| **CTL2A** | Expressed in activated T-cells | 4/4 (100%) | 0/5 (0%) | Strong |
| **PGS2** | Involved in collagen fibril formation; belongs to damage-associated molecular pattern (DAMP) family | 4/4 (100%) | 0/5 (0%) | Strong |
| **BIP** | Molecular chaperone, protein quality control, BBB disruption | 4/4 (100%) | 1/5 (20%) | Strong |
| **GFAP** | Astrocyte activation marker | 3/4 (75%) | 0/5 (0%) | Strong |
| **LAG3** | Immune checkpoint protein | 3/4 (75%) | 0/5 (0%) | Strong |
| **NPTX1** | Synaptic plasticity regulator | 3/4 (75%) | 0/5 (0%) | Strong |
| **CHI3L1** | Astrocyte-derived inflammatory marker | 3/4 (75%) | 1/5 (20%) | Strong |
| **ACTA** | Pericyte contractile protein | 3/4 (75%) | 1/5 (20%) | Strong |
| **LAMB2** | Basement membrane component | 3/4 (75%) | 1/5 (20%) | Strong |
| **CSPG5** | Neural development | 4/4 (100%) | 1/5 (20%) | Strong |
| **NFASC** | Axon-glia interactions | 4/4 (100%) | 1/5 (20%) | Strong |
| **RELN** | Synaptic plasticity, neuronal migration | 4/4 (100%) | 1/5 (20%) | Strong |
| **C1QA** | Complement activation | 2/4 (50%) | 0/5 (0%) | Moderate |
| **VCAM1** | Endothelial adhesion molecule | 2/4 (50%) | 0/5 (0%) | Moderate |
| **CADH5** | VE-cadherin, endothelial junctions | 2/4 (50%) | 1/5 (20%) | Moderate |
| **CX3CL1** | Microglial chemokine | 2/4 (50%) | 0/5 (0%) | Moderate |
| **UCHL1** | Neuronal injury marker | 2/4 (50%) | 1/5 (20%) | Moderate |

**ACTA** – $\alpha$ smooth Muscle Actin (Actin, aortic smooth muscle); **BIP** - Binding Immunoglobulin Protein (also known as GRP78/HSPA5); **C1QA** - Complement C1q A Chain; **CADH5** - Cadherin 5 (VE-Cadherin); **CHI3L1** - Chitinase 3 Like 1 (YKL-40); **CTL2A** - Protein CTLA-2- $\alpha$ ; **CSPG5** - Chondroitin Sulfate Proteoglycan 5 (Neuroglycan C); **CX3CL1** - C-X3-C Motif Chemokine Ligand 1 (Fractalkine); **GFAP** - Glial Fibrillary Acidic Protein; **LAG3** - Lymphocyte Activation Gene 3; **LAMB2** - Laminin Subunit Beta 2; **MMP2** - Matrix Metalloproteinase-2; **NFASC** - Neurofascin; **NPTX1** - Neuronal Pentraxin 1; **PGS2** - Decorin; **RELN** - Reelin; **UCHL1** - Ubiquitin C-Terminal Hydrolase L1; **VCAM1** - Vascular Cell Adhesion Molecule 1.

**Table 2:** Non-dep-preferred proteins.

| Protein | Function | Non-dep Detection | Dep Detection | Evidence Strength |
| --- | --- | --- | --- | --- |
| **TAGL3** | Cytoskeletal regulation | 4/5 (80%) | 0/4 (0%) | Strong |
| **KV5AB** | Kappa light chain variable region | 4/5 (80%) | 1/4 (25%) | Strong |
| **CALB1** | Calcium-binding neuroprotection | 4/5 (80%) | 1/4 (25%) | Strong |
| **LV1C** | Lambda light chain variable region | 3/5 (60%) | 0/4 (0%) | Moderate |
| **SUMO2/3** | Protein modification, stress response | 3/5 (60%) | 1/4 (25%) | Moderate |

**CALB1** - Calbindin 1; **KV5AB** - Ig kappa chain V-V region HP R16.7; **LV1C** - Ig lambda-1 chain V region S178; **SUMO2/3** - Small Ubiquitin-like Modifier 2/3; **TAGL3** - Transgelin-3.
\* Note: Proteins listed represent minimum estimates of group differences due to limited power for detecting moderate effect sizes.

The proteomic signatures identified here represent minimum estimates of alcohol dependence-associated changes, as our study design provided adequate power only for detecting large effect sizes. This suggests that the true extent of proteomic alterations in alcohol dependence may be substantially greater than reported.

### Non-dep-Specific CSF Proteins Indicate Preservation of Protective Mechanisms

Proteins unique to the Non-dep ethanol group revealed several protective and regulatory mechanisms that appear to be lost in alcohol dependence. CALB1, a critical calcium-binding protein primarily expressed in inhibitory neurons, was preferentially detected in Non-dep (1/4 Dep vs 4/5 Non-dep). In a *Calb1^−/−^*KO mouse model, *Calb1* was found to regulate sex-specific behavioral differences. KO was related to reduced anxiety in both sexes; however, fear, memory and social behavior changes were observed in male but not female mice ^18^. The same study also found an effect on histone deacetylase transcript levels (*Hdac3* and *Hdac4*) in the amygdala and prefrontal cortex (PFC) between WT and KO animals, suggesting a role for CALB1 in epigenetic regulation within these brain regions ^18^. CALB1 has also garnered interest for its neuroprotective properties in neurodegenerative conditions such as Alzheimer’s ^19^ and Parkinson’s Disease ^20^. Preserved calcium homeostasis proteins (CALB1; Calretinin-2, CALB2) in Non-dep mice suggest that neuronal calcium buffering capacity is maintained, which may protect against excitotoxicity and cell death ^21–23^. The loss of these protective mechanisms in dependence could predispose neurons to calcium-mediated damage.

The preferential detection of numerous proteins associated with cellular stress response and protein degradation in Non-dep animals suggests that protein modification systems crucial for cellular stress responses are maintained. SUMO2/SUMO3 conjugation proteins (1/4 Dep vs 3/5 Non-dep) are associated with neuroprotective mechanisms ^24^, and it has been suggested that they act through multiple mechanisms to carry out protein quality control in the brain. The downregulation of Transgelin-3 (TAGL3; 0/4 Dep vs 4/5 Non-dep), a regulator of cytoskeletal regulation and smooth muscle function, has been implicated as a driver of neuroinflammation in astrocytes ^25^. Multiple specific immunoglobulin variable regions were also preferentially detected in the Non-dep group, suggesting that adaptive immune surveillance mechanisms may be compromised in alcohol dependence. These included kappa light chain variable regions (KV5AB cluster: 1/4 Dep vs 4/5 Non-dep; KV5A7: 0/4 Dep vs 2/5 Non-dep), lambda light chain variable regions (LV1C: 0/4 Dep vs 3/5 Non-dep), and heavy chain variable regions (HVM02/03 and HVM57: 0/4 Dep vs 2/5 Non-dep each). The detection of these antibody components in the Non-dep group may be associated with immunosuppression and reduced adaptive immune capacity in alcohol dependence ^26^.

### Blood-Brain Barrier Disruption in Alcohol Dependence

The BBB is a layer of tightly packed semi-permeable cells that act as gatekeepers to the CNS; the BBB selectively allows or disallows entry of molecules from systemic circulation into the neural compartment. Chronic alcohol exposure can disrupt the BBB, leading to increased neuroinflammation and cognitive decline ^27,28^. A significant finding in this study was the detection of MMP2 in all Dep mice but absence in the Non-dep group. MMP2 is an enzyme critically involved in ECM degradation and BBB breakdown, making its exclusive presence in dependent mice suggestive of vascular compromise ^29^. The endoplasmic reticulum chaperone BiP (BIP; gene = *Grp78*/*Hspa5*) also displayed a strong Dep preference, with detection in all animals in the Dep group and 1/5 Non-dep mice. BIP was found to be associated with both endoplasmic reticulum dysfunction and the disruption of endothelial tight junctions *in vitro*, as its abundance was upregulated following ethanol exposure in a rat brain endothelial cell line, RBE4, and coincided with increased oxidative stress ^30^. In humans, elevated levels of MMP2 can be detected in blood serum of individuals with alcoholic liver disease and was found to correlate with the development of liver cirrhosis ^31^. Similarly, increased BIP levels were found to be positively correlated with alcohol consumption grade and reflected the degree of alcohol-induced damage in human liver biopsy samples ^32^. A less pronounced but consistent pattern could be observed with the tendency toward Dep-detection for endothelial junction proteins including VE-cadherin (CADH5; 2/4 Dep vs 1/5 Non-dep) and vascular cell adhesion molecule-1 (VCAM1; 2/4 Dep vs 0/5 Non-dep), which may indicate early signs of cellular response to vascular endothelial damage and compromised BBB function ^33,34^.

Evidence for pericyte involvement was demonstrated by the Dep-preferred detection of alpha-smooth muscle actin (ACTA) in 3/4 dependent mice vs only 1/5 Non-dep mice. ACTA is a major contractile protein normally confined within pericytes ^35^, and its presence in CSF suggests pericyte detachment or death, a hallmark feature of BBB dysfunction. This finding suggests that alcohol dependence involves the pericyte component of the neurovascular system responsible for BBB regulation.

The ethanol dependent-preferred detection of laminin beta-2 (LAMB2) in 3/4 dependent animals versus only 1/5 Non-dep is suggestive of compromised basement membrane integrity in alcohol-dependent mice and suggests structural damage to the neurovascular unit ^36^. The detection of fibulin-4 (FBLN4) in all dependent mice but only 1/5 Non-dep, and fibulin-5 (FBLN5) in 3/4 Dep mice but no Non-dep, could indicate leakage of these vascular scaffolding proteins from the ECM. Since FBLN4 and FBLN5 are essential for maintaining blood vessel strength and elasticity ^37,38^, their presence in CSF suggests vascular matrix degradation and compromised vessel integrity in alcohol dependence.

### Neuroinflammation

The presence of several key markers in the alcohol dependent group is indicative of dependence-associated neuroinflammation. GFAP, the gold-standard biomarker for astrocyte activation, was detected in 3/4 Dep animals but 0/5 Non-dep, representing one of the strongest group differences observed. GFAP has been extensively validated as a biomarker for traumatic brain injury and neuroinflammation ^39^, and cortical astrocyte activation has been linked to the regulation of ethanol consumption in mice ^40^. These findings highlight the potential of GFAP as an indicator of chronic excessive alcohol exposure that induces dependence or related neurological damage.

Similarly, Chitinase-3-like protein 1 (CHI3L1) showed ethanol dependence preference (3/4 Dep vs 1/5 Non-dep). CHI3L1 is primarily secreted by activated astrocytes and has been established as a reliable biomarker for inflammatory CNS conditions. CHI3L1 levels in CSF have been associated with neuroinflammation across multiple neurological diseases, including multiple sclerosis and Alzheimer’s disease ^41^. The preferential detection of CHI3L1 in dependent mice is suggestive of neuroinflammation; however, further studies are required to validate its association with the transition to alcohol dependence.

The chemokine CX3CL1 (fractalkine), which mediates microglial activation and neuroinflammation ^42^, was also Dep-preferred (2/4 Dep vs 0/5 Non-dep). We detected other neuroinflammation-related proteins, including the immune checkpoint protein and regulator of cytokine production, lymphocyte activation gene 3 (LAG3), which was detected in 3/4 Dep animals and not detected in Non-dep. One study noted that an increase in soluble and membranous forms of LAG3 were observed in primary cultured microglia *in vitro* in response to interferon-γ (IFN-γ) treatment, with additional *in vivo* experiments confirming that LAG3 levels increase occurs via the IFN-γ/STAT1 pathway ^43^. LAG3 detection in dependent animals is consistent with the neuroinflammatory profile in the Dep group and supports the hypothesis that immune system dysregulation is driven by chronic alcohol consumption.

### Alcohol-Induced Changes to the CSF Proteome

The ethanol-dependence-specific proteins indicate a molecular response involving vascular damage, neuroinflammation, and cellular stress. The presence of VE-cadherin in CSF is particularly noteworthy, as this protein is normally tightly localized to endothelial junctions ^44^, therefore its detection in the Dep group suggests BBB compromise. This finding aligns with recent mechanistic studies showing that alcohol disrupts VE-cadherin expression and BBB integrity through multiple pathways ^27^.

GFAP, CHI3L1, and CX3CL1 detection in dependent animals is consistent with a neuroinflammation signature suggestive of glial activation and CNS distress. The ability to detect these proteins within 4-5 weeks of dependence initiation offers critical insight into the rapid neurological decline that occurs following sustained alcohol exposure.

### Immune System Dysregulation and Loss of Protective Mechanisms

STRING network analysis of non-Dep-preferred proteins revealed maintained physiological homeostasis mechanisms that appear to be down-regulated in alcohol dependence (Figure 3a). The detection of proteins involved in acute phase response (CRP), and lipid transport (APOB) suggests preserved regulatory capacity in moderate alcohol-exposed Non-dep mice.

**Figure 3:**
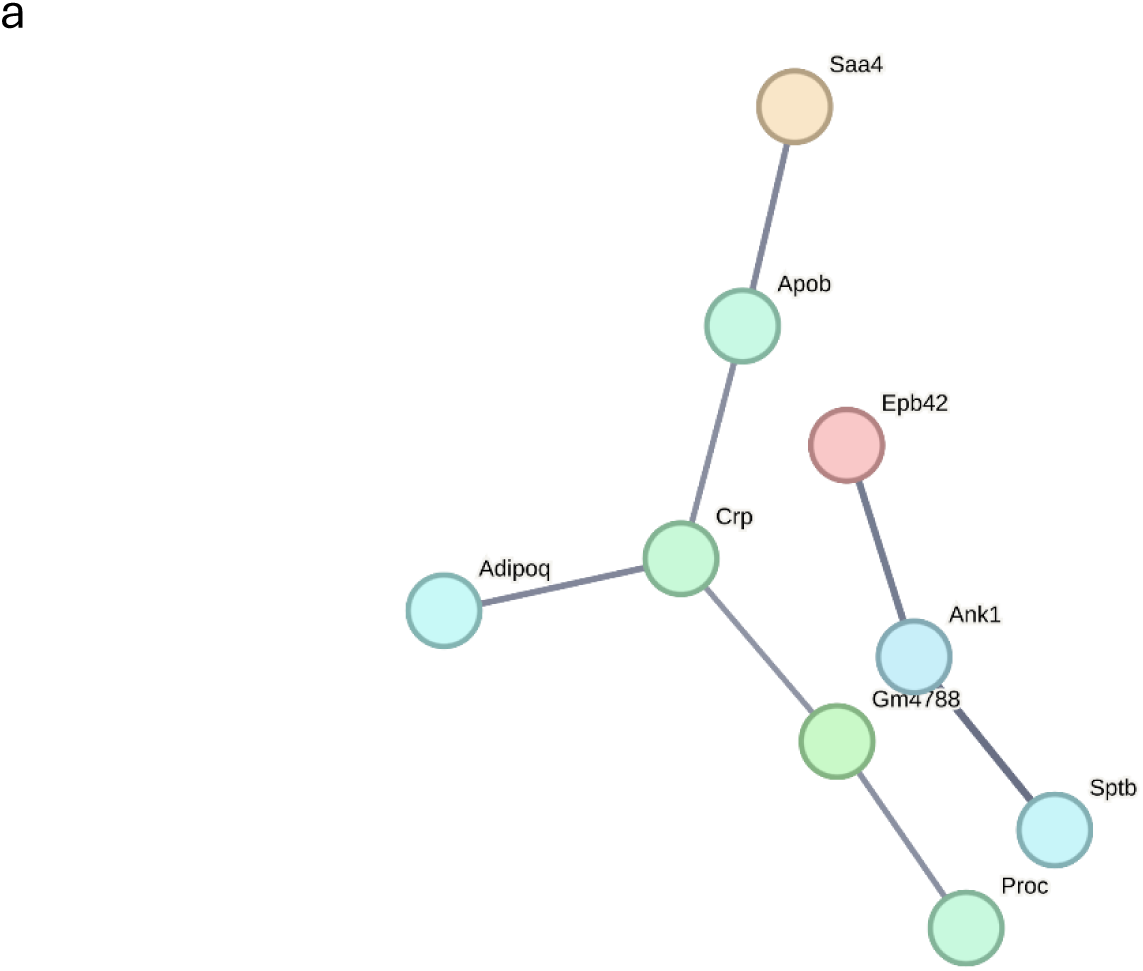
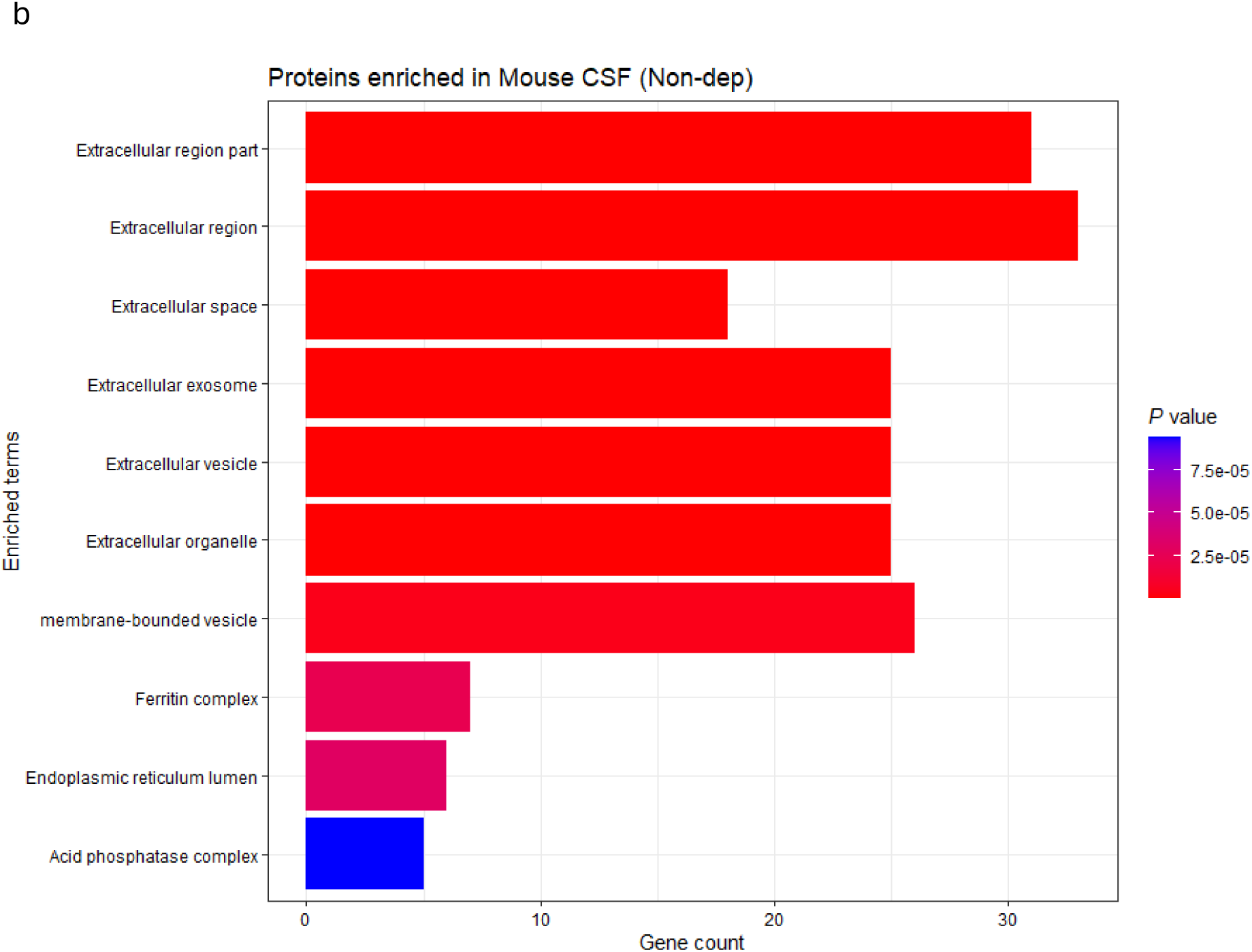
a) STRING protein-protein interaction (PPI) network analysis of proteins uniquely identified in alcohol-exposed, Non-dependent mice (Non-dep). Colored nodes: query proteins and first shell of interactors. Protein interactions were filtered at high confidence (≥0.700; line width is positively correlated with confidence level) and unconnected nodes removed. b) EnrichR analysis against Jensen COMPARTMENTS database shows enrichment of extracellular compartments among Non-dep-preferred proteins. Bar length represents gene count; color intensity indicates statistical significance (*P*-values).

Analysis of Non-dep-preferred proteins against the Jensen COMPARTMENTS database revealed a markedly different pattern than Dep-specific proteins, suggesting maintained cellular organization and protective mechanisms in alcohol-exposed but Non-dep mice (Figure 3b). While extracellular region proteins were enriched (∼31 proteins), the overall extracellular protein diversity was substantially lower than observed in dependent mice (∼95 proteins in the Dep group), suggesting dependent mice have more uncontrolled protein release. Importantly, Non-dep-specific proteins showed significant enrichment for protective complexes, including the ferritin complex, reflecting maintained iron homeostasis. The presence of endoplasmic reticulum lumen proteins suggests preserved cellular organization and protein processing capacity. This compartment profile supports the interpretation that alcohol-exposed Non-dep mice maintain protective mechanisms and cellular integrity, as opposed to the extensive extracellular protein leakage observed in dependent mice. The controlled extracellular protein presence in the Non-dep group likely represents normal physiological protein turnover rather than pathological cellular compromise.

In contrast, alcohol-dependent mice showed enhanced classical pathway activation, evidenced by the presence of C1QA and C6 proteins specifically in the Dep group (Figure 4a). This represents dysregulated innate immune activation, while suppressed adaptive immunity is also evidenced in the same animals. C1QA activation of the classical complement pathway and C6 participation in membrane attack complex formation indicate both initiation and terminal complement activation, which are known contributors to neuroinflammation and synaptic pruning. Complement regulatory proteins (CFHR1, CFHR4) detected in the Non-dep group suggest that complement control is maintained ^45^, which is juxtaposed with the complement activation (C1QA, C6) seen in dependent mice. This shift from regulated to dysregulated complement activation may contribute to neuroinflammation and tissue damage observed in models of alcohol dependence ^46–48^.

**Figure 4:**
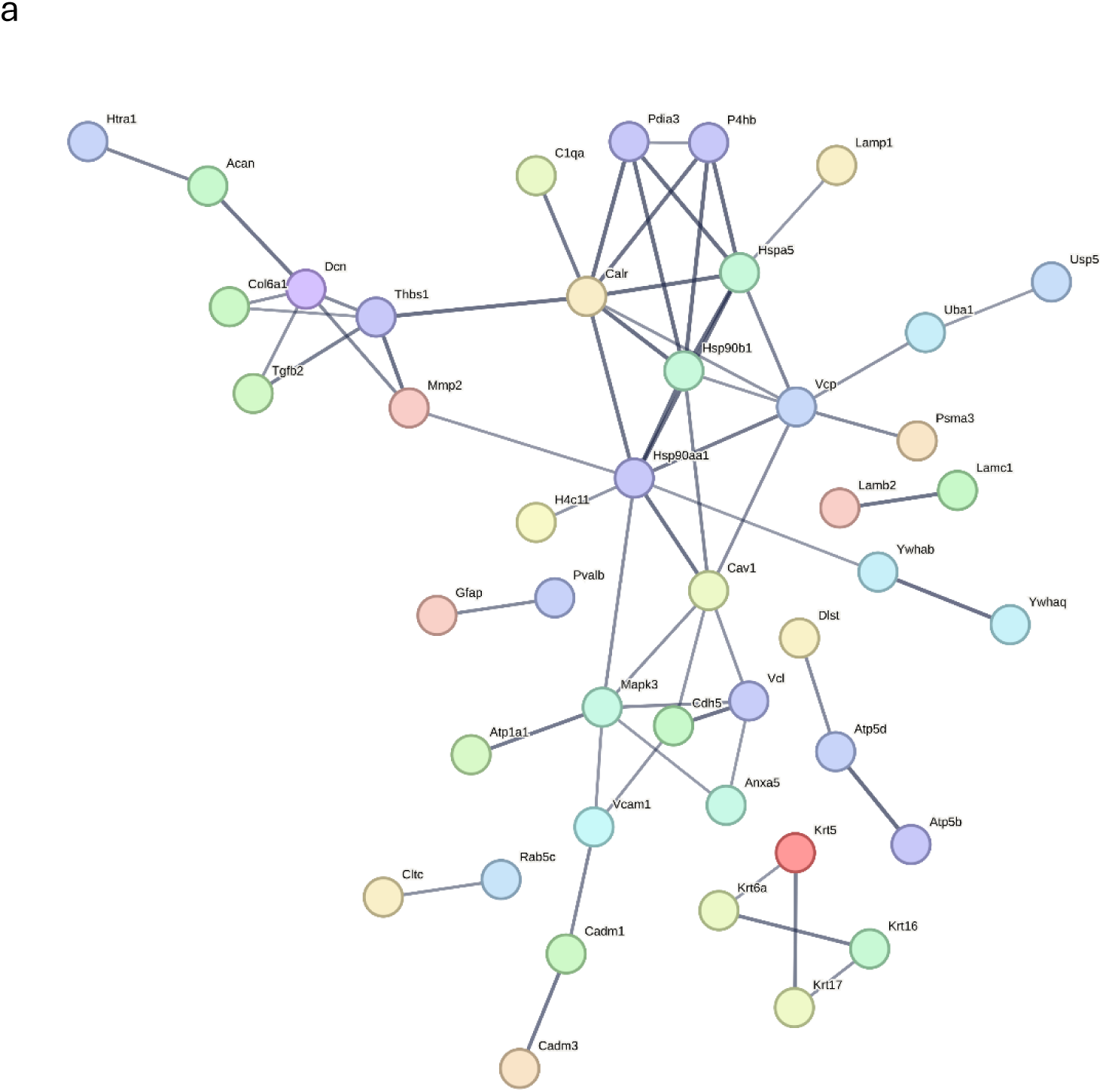
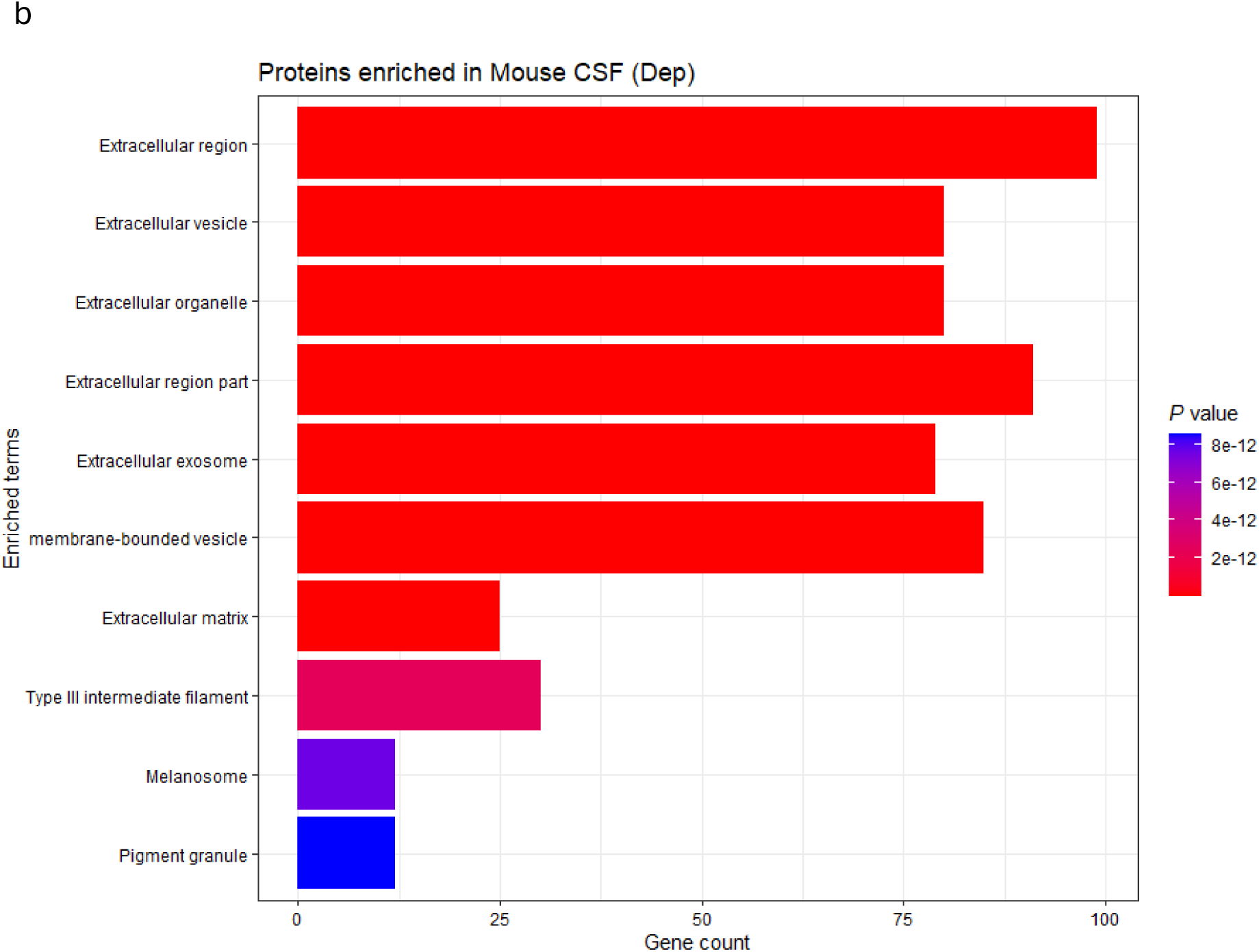
a) STRING protein-protein interaction (PPI) network analysis of proteins identified in alcohol-dependent mice (Dep). Colored nodes represent query proteins and their first-shell interactors. Protein interactions were filtered for high confidence (≥0.700; line width correlates with confidence level), and unconnected nodes were removed. PPIs between complement family member C1QA, heat shock proteins HSPA5 and HSP90B1, as well as proteasome and protein degradation proteins PSMA3, UBA1, and USP5, were scored as high to extremely high confidence (≥0.700 – 0.900). b) EnrichR analysis against the Jensen COMPARTMENTS database reveals significant enrichment of extracellular compartment proteins among Dep-preferred proteins. Bar length indicates gene count; color intensity reflects statistical significance (*P*-values).

### Cellular Stress and Protein Quality Control

Multiple indicators of cellular stress were found only in Dep mice. Heat shock proteins (HSPA5, HSP90B1) and protein disulfide isomerases (P4HB, PDIA3) suggest endoplasmic reticulum stress responses (Figure 4a). Proteasome subunit alpha-3 (PSMA3) and ubiquitin-related enzymes (UCHL1, UBA1, USP5) show activation of the protein degradation system. These proteins indicate upregulation of the cellular stress response and attempts to handle cell and tissue damage but also links to misfolded protein accumulation and aggregation as proposed by others ^49,50^. Metabolic disturbance was evidenced by the presence of multiple ATP synthesis enzymes (ATP5B, ATP5D) and glycolytic enzymes (PFKAM), suggesting mitochondrial dysfunction and energy metabolism disruption.

Overall, cellular compartment enrichment analysis provided evidence for structural compromise in the alcohol-dependent mice (Figure 4b). The Dep-preferred proteins showed significant enrichment for extracellular compartments. This substantial extracellular protein presence, encompassing proteins associated with extracellular vesicles (EVs), organelles, and exosomes, suggests that alcohol dependence promotes increased leakage of cellular proteins into CSF while also promoting intercellular communication via EVs, which are known mediators of neuroinflammation in various neurological conditions ^51,52^. The enrichment of structural components, including Type III intermediate filaments, indicates that cellular integrity may be compromised, with cytoskeletal and membrane proteins detected in the extracellular space. These findings support the interpretation that in this model, alcohol dependence involves significant BBB dysfunction and cellular compromise, where normal compartmentalization between cellular and extracellular spaces becomes disrupted. The enhanced proteasome activity and cellular stress responses suggest that alcohol dependence overwhelms normal protein homeostasis mechanisms.

### Functional Pathway Analysis

KEGG (Kyoto Encyclopedia of Genes and Genomes) pathway enrichment analysis of Dep-preferred proteins revealed significant enrichment of pathways consistent with the observed proteomic signatures (Figure 5). The most significantly enriched pathway by gene count was PI3K-Akt signaling pathway (FDR = 3.5×10⁻^4^), indicating dysregulated cell survival and proliferation signaling that may contribute to altered neuronal responses to alcohol exposure. Prion disease pathway enrichment (FDR = 7.9×10⁻^4^) suggests protein misfolding and aggregation processes, consistent with the detected cellular stress response proteins. Fluid shear stress and atherosclerosis pathway involvement (FDR = 7.0×10⁻^5^) indicates vascular dysfunction patterns typically associated with cardiovascular disease, directly supporting the observed BBB breakdown evidenced by MMP2, VE-cadherin, and pericyte damage markers. Protein processing in endoplasmic reticulum pathway enrichment (FDR = 1.5×10⁻^4^) highlights the cellular stress response signatures, including the detection of heat shock proteins and protein quality control markers. Focal adhesion pathway involvement (FDR = 3.5×10⁻^4^) aligns with the detected extracellular matrix degradation and pericyte detachment, while estrogen signaling pathway enrichment (FDR = 7.0×10⁻^5^) may reflect loss of neuroprotective mechanisms. Collectively, these pathway enrichments suggest that alcohol dependence is characterized by dysregulated cellular signaling, protein homeostasis failure, vascular compromise, and loss of protective mechanisms.

**Figure 5:**
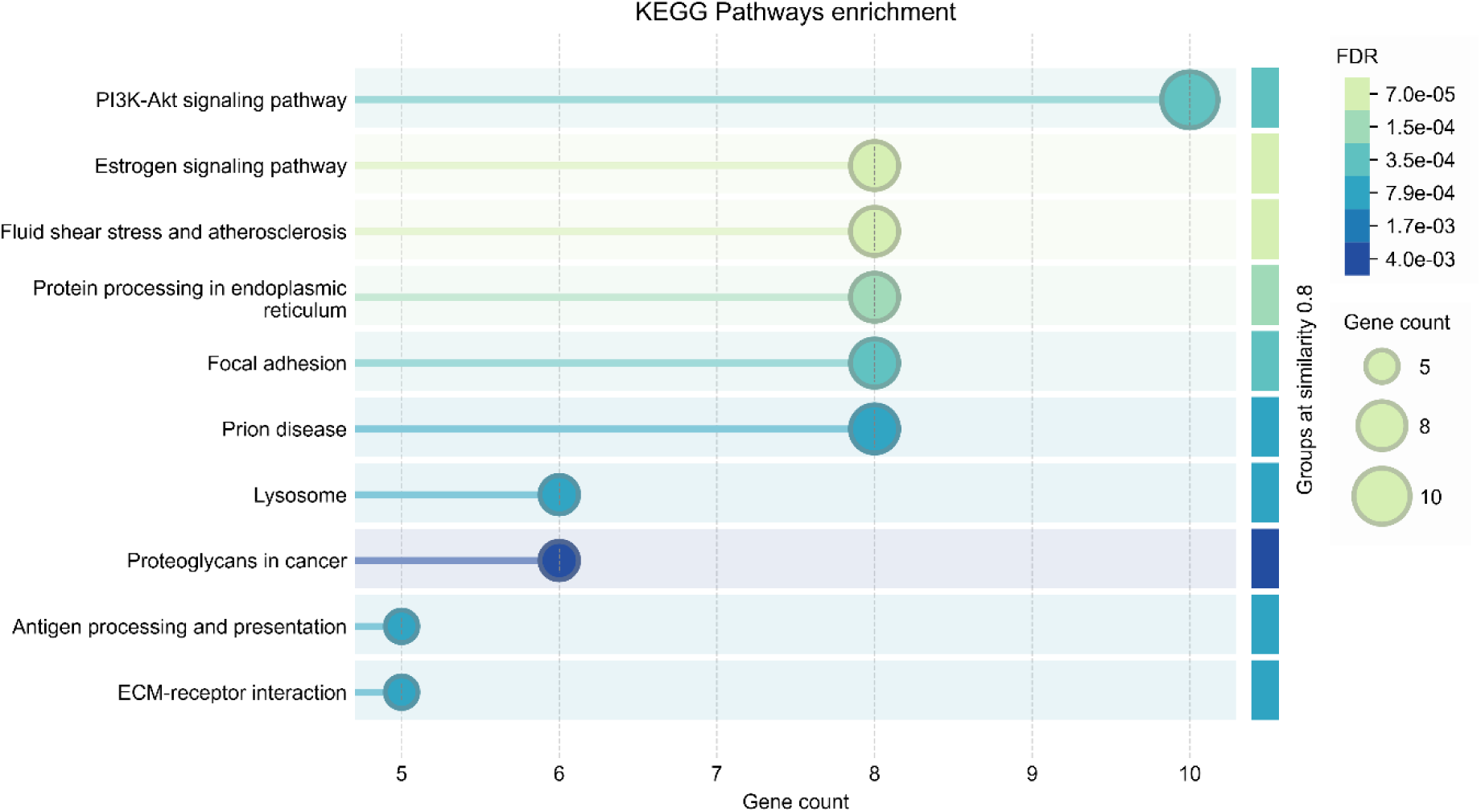
KEGG pathway enrichment analysis of Dep-specific proteins reveals multi-system dysfunction in alcohol dependence. Functional enrichment analysis was performed using STRING database on proteins detected in ≥2/4 alcohol-dependent and ≤1/5 alcohol-exposed Non-dep mice. Circle size represents gene count; color intensity indicates statistical significance (FDR values).

### Synaptic and Neuronal Dysfunction

Several proteins involved in synaptic function and neuronal integrity showed strong, but not exclusive, Dep preference. Neurofascin (NFASC), crucial for axon-glia interactions and node of Ranvier formation ^47^, was detected in all Dep animals and 1/5 Non-dep (4/4 vs 1/5). RELN, critical for synaptic plasticity and neuronal migration ^53^, showed identical preferential detection in dependent mice (4/4 vs 1/5).

Chondroitin sulfate proteoglycan 5 (CSPG5) was also Dep-preferred (4/4 vs 1/5). CSPG5 is involved in neural development and synaptic regulation ^54^, and was found to be primarily expressed in GFAP+ radial glial cells in a transgenic mouse model focusing on neural stem cells and neural maturation ^55^. Neuronal pentraxin 1 (NPTX1) was detected in 3/4 Dep animals but absent in Non-dep. NPTX1 serves dual roles as both a proapoptotic protein induced by reduced neuronal activity and a regulator of Kv7.2 potassium channel surface expression, which modulates neuronal excitability ^56^. NPTX1 is also known for its role in regulation of AMPA receptor clustering and synaptic strength ^57^. Taken together, these findings suggest that alcohol dependence disrupts synaptic organization and axon-glia communication.

### Limitations

The presence of multiple keratin proteins (K2C5, K1C10, K2C1B) and hemoglobin (Hb) in the Dep-specific protein list raises questions about potential sample contamination or tissue damage. However, the biological pattern of neuroinflammation, BBB disruption, and synaptic dysfunction supports the biological significance of these findings rather than pure technical artifacts. We reviewed relevant literature that may shed light on the identification of these proteins by proteomics in mouse CSF and found one recent methodological study specifically aimed at CSF collection in murine models for the purpose of sample purity and contaminant mitigation ^58^. Interestingly, even though that study ^58^ reported undetectable levels of Hb in CSF samples with ELISA, proteomics analysis identified Hb, in addition to multiple classes of keratins, which were analyzed using an earlier version of the same DIA-NN pipeline (results available from ProteomeXchange, dataset identifier: PXD053568) ^58^. Overall, the number of protein IDs were similar between our study and this comparator methodological study. This demonstrates that MS-based proteomics can detect trace amounts of these proteins below the detection limits of conventional immunoassays as well as supporting the legitimacy of these identifications outside of contamination. In the current study, the CSF proteome complexity was similar across all samples (Figure 1a), which would likely not be the case if contamination had contributed significantly to the proteomic findings in the Dep group. Given that numerous proteins indicate BBB leakage and vascular damage, further studies should focus on confirming these findings and any associated processes which could lead to cell and tissue breakdown or leakage into CSF. Increasing the sample size in future studies will allow for a deeper evaluation and interpretation of dependence-associated alterations in the neurological compartment.

The use of a stringent detection threshold (≥2/4 animals in one group, ≤1/5 in the comparison group) was employed to minimize false discoveries while maintaining sensitivity for biologically relevant changes. This approach identified 140 Dep-preferred and 67 Non-dep-preferred proteins, providing high-confidence biomarker candidates for validation studies. Post-hoc power analysis revealed that our study design (*n* = 4-5 per group) provided >99% power to detect large effect sizes (detection rates of 100% vs 0%) but only 38% power for moderate effects (75% vs 20% detection rates). This suggests that our findings likely underestimate the true number of proteins differing between groups, particularly those with moderate effect sizes. The limited *n* value in the current study underlines the necessity of performing a follow up validation study with a larger sample size, calculated to be *n* = 12-15/group for 0.8 power for the detection of moderate effect sizes. This would also enable quantitative comparison between shared proteins, thus providing a more comprehensive analysis and understanding of alterations to the CSF proteome in AUD models.

Another limitation is that this cohort of mice was chronically administered IL-6R which might have attenuated some general IL-6 related mechanisms. However, importantly, the effects seen in the Dep group were not ameliorated by IL-6R treatment, suggesting that there are multiple pathways contributing to the dependence-associated neuroinflammation reported here. To further investigate whether the IL-6R antibody was detectable in the mouse CSF samples, we searched the proteomics data against the *Rattus norvegicus* proteome and filtered the output to include peptides from the IGG2B protein (see Supplementary file 2 for the list of identified sequences filtered at 1% FDR). Out of three identified peptide sequences belonging to IGG2B, one sequence was unique to rat (DILLISQNAK), which was identified in all CSF samples. Others have shown that intravenously administered antibodies can cross the BBB even when it is not compromised ^59^, and IgG antibody has been detected in the interstitial brain fluid in naïve mice by other methods ^60,61^. This presents MS-based proteomics as a potential screening tool for detecting antibodies in the CNS following their peripheral applications in small volumes of CSF.

While the current study lacked the statistical power to perform sex stratification, expanding this work to identify sex-specific differences in dependence-mediated molecular mechanisms of BBB dysfunction and immune dysregulation is warranted. Further considerations or expansions in this area of research may also include changes in abundance of histone deacetylases and are known to be affected in addiction models ^18^. This suggests that the transient and longer-term molecular mechanisms of addiction may be potentiated by epigenetic dynamics and is an ongoing area of research in the field of addiction ^62–65^. Future studies may focus specifically on histone post-translational modification (PTM) analysis and alterations to normal PTM patterns in the transition to alcohol dependence ^66^.

## Conclusions

This study identified differences in the CSF proteomes between alcohol dep and non-dependent moderate alcohol drinking mice indicative of a shift from maintained protective mechanisms to neuroinflammation and loss of immune regulation in alcohol dependence. Proteins detected in the Non-dependent group support active protective processes, including complement regulation, anti-inflammatory signaling, and neuronal calcium homeostasis, which are lost or diminished during the transition to dependence. In contrast, the alcohol dependent-specific proteins indicate multiple pathological responses involving BBB breakdown, neuroinflammation, and cellular stress. The contrast between the two proteomes suggests that alcohol dependence involves a combination of pathological processes and loss of protective mechanisms. The detection of proteins involved in the neurovascular system such as endothelial cells (VE-cadherin), pericytes (ACTA), and basement membrane (LAMB2, fibulins) suggests that alcohol dependence affects all components of the BBB architecture. Importantly, we provide a link to human studies that have detected two of the strong candidates identified here, MMP2 and BIP, in human serum and liver samples ^31,32^, indicating that these putative markers of alcohol dependence detected in CSF may not be confined to the central nervous system. This presents an opportunity to explore alternative diagnostic routes of alcohol-dependence-related CNS damage via peripheral blood ^67^ or tissue sampling in future studies.

Notably these proteomic data are in agreement with our recent studies (using the same CIE-2BC mouse model) showing that alcohol dependence causes a strong neuroimmune response in astrocytes in the nucleus accumbens, including activation of interferon and interleukin pathways, while moderate drinking alone caused homeostatic responses in astrocytes but did not show evidence of a neuroimmune activation ^68^. While these findings provide preliminary evidence for distinct proteomic signatures in alcohol dependence, the identified proteins represent minimum estimates due to limited statistical power for moderate effect sizes. The identification of clinically validated biomarkers (GFAP, UCHL1, CHI3L1) in the dependence group suggests potential for developing diagnostic and monitoring tools for alcohol-related brain injury. Furthermore, the comprehensive nature of the pathological response, combined with the loss of protective mechanisms, indicates that alcohol dependence creates a vulnerable brain state that may predispose to accelerated aging and neurodegenerative processes. Validation in larger cohorts is essential to capture the full scope of dependence-associated molecular changes.

## Methods

### Animals

The male and female C57BL/6J mice (*n* = 4 and 5; 29.2±1.63 g and 22.4±0.55 g, respectively) obtained from The Jackson Laboratory (Bar Harbor, ME) were housed in a temperature-and humidity-controlled room (12 h reverse light cycle) and provided with food and water *ad libitum*. All the animal procedures comply with the ARRIVE guidelines and were approved by The Scripps Research Institute (TSRI) Institutional Animal Care and Use Committee (IACUC #09-0006), consistent with the National Institutes of Health Guide for the Care and Use of Laboratory Animals.

### Chronic intermittent ethanol – two bottle choice model

The mice used in this study were part of a cohort consisting of a total of 12 males and 12 females undergoing the chronic intermittent ethanol vapor-2 bottle choice paradigm (CIE-2BC) as described ^9,10,69,70^ and treatment with the InVivoMAb anti-mouse IL-6R antibody (#BE0047, BioXCell).

To acclimate the mice to ethanol, they were singly housed and allowed 24 h access to 15% ethanol in addition to their normal water and food for 4 days (Mon-Fri). After this, mice were returned to their original group housing. For an additional 15 days (5 days per week for 3 weeks) mice were singly housed for two hours with access to two drinking tubes, one containing 15% ethanol and the other containing water (i.e. two bottle choice or 2BC) for 30 min before the lights were turned off. After this baseline drinking, the mice were assigned to the ethanol Non-dep or ethanol-dependent groups, with groups matched for baseline ethanol and water consumption. The ethanol Non-dep mice had a moderate level of voluntary ethanol consumption through limited access 2BC drinking sessions. In contrast, the ethanol-dependent mice received intermittent vapor exposure in addition to 2BC drinking sessions, resulting in the escalated voluntary ethanol intake. The mice underwent 6 cycles of 4 days of CIE (16 h ethanol vapor/8 h air) with their home cages inserted into vapor chambers (La Jolla Alcohol Research, La Jolla, CA) and 3 days forced abstinence followed by 5 days of 2BC (same parameters as in baseline training) and 2 days forced abstinence (Figure 6). The Non-dep mice were exposed only to air in their home cages. Immediately before each ethanol vapor/air exposure, the dependent mice were injected (i.p.) with 1.75 g/kg ethanol + 68.1 mg/kg pyrazole (Sigma, St Louis, MO) and Non-dep mice with 68.1 mg/kg pyrazole. The blood ethanol levels (BELs) were determined weekly using an Agilent 7820A GC coupled to a 7697A (headspace-flame-ionization) and were compared with and calibrated using a 6-point serial diluted calibration curve of 300 mg/dl ethanol (Cerilliant E-033). Results were used to adjust ethanol drip rates in the vapor chambers to achieve BELs of 150-250 mg/dL.

**Figure 6:**
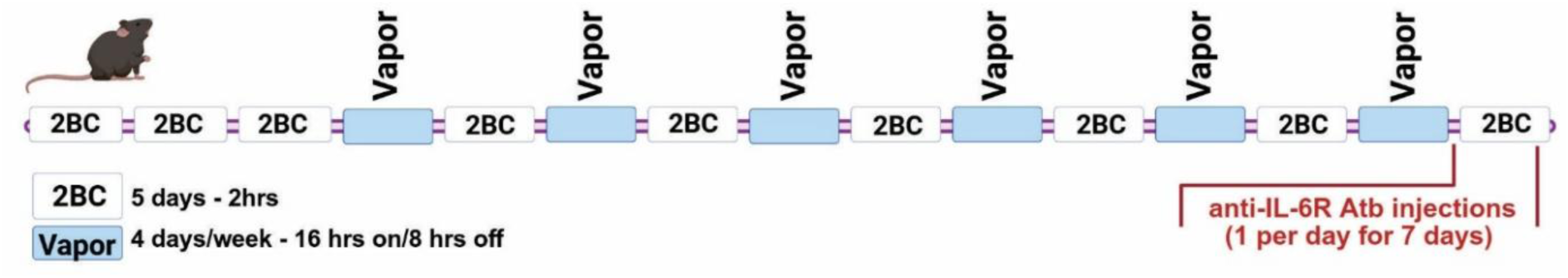
Schematic of the 2BC-CIE paradigm.

The mice used in this study were injected with the InVivoMAb anti-mouse IL-6R antibody (BE0047, BioXCell) intraperitoneally (200 microgram/day) for 7 consecutive days. The first injection of the antibodies was 24 h after removal from the last vapor exposure. Thus, the antibody injections span over the forced WD after the vapor session and 4 days of a 2BC testing period. All injections were conducted to match the injection time during the 2BC testing period, 30 minutes before the drinking session. On the final day of antibody exposure CSF was collected between 2 and 4 h post injection.

### CSF collection

After heavily anesthetizing the mice with isoflurane, the mice were secured in a stereotaxic unit, which continued to administer isoflurane. The area above the cisterna magna (located between the cerebellum and the dorsal surface of the medulla oblongata and receiving CSF from the 4th ventricle) was then isolated by cutting and deflecting the layers of skin and muscle at the back of the head. A pulled pipette was used to collect the cerebrospinal fluid by capillary pressure. Approximately 5 −20μl of cerebrospinal fluid was collected over no more than 30 min. and the mice were immediately euthanized. The collected CSF samples were stored at – 80 °C until analyzed.

### Proteomics Sample Preparation

Mouse CSF samples were processed for proteomic analysis using a modified filter-aided sample preparation protocol, as previously described ^71^. Briefly, samples (∼5 – 15 µL) were solubilized in 20 µL of lysis buffer containing 1% w/v sodium deoxycholate, 1 % w/v sodium dodecyl sulphate, 100 mM dithiothreitol (DTT) and 100 mM Tris-HCl pH 8.5, supplemented with protease inhibitors (Pierce Protease Inhibitor mini tablets, Thermo Scientific). Samples in lysis buffer were vortexed, heated to 95 °C for 3 min, and sonicated in a water bath for 5 min. Samples were incubated on ice for 10 min before being loaded onto Nanosep 30K centrifugal filter devices (Cytiva) and centrifuged for 15 min at 14,000 x*g* and 21 °C until all the sample had passed through the filter. Samples were reduced with 25 mM DTT in 8 M urea for 1 h at RT, then centrifuged as in the previous step. Filters were washed once with 8 M urea, then 50 mM iodoacetamide in 8 M urea was added to the filters and incubated in the dark at RT for 20 min. Filters were centrifuged once and washed again with 8 M urea, followed by another wash with 100 mM ammonium bicarbonate pH 8-8.5 (AMBIC). Trypsin suspended in 100 mM AMBIC was added to each filter and an additional 60 µL of 100 mM AMBIC was added to each filter. The filter lids were left open, and samples incubated in a humidified chamber overnight at 37 °C. The next day, filters were transferred to clean 1.5 mL protein lobind tubes (Eppendorf) and centrifuged to elute peptides. One additional elution was performed by adding another 40 µL 100 mM AMBIC to the filters and repeating the centrifugation. Samples were acidified with formic acid (FA) to a final concentration of 0.1%. A peptide assay was performed to determine peptide concentration following the manufacturer’s instructions (Pierce Colorimetric Quantitative Peptide Assay, Thermo Scientific). A volume equivalent to 350 ng peptides was loaded onto Evotips (Evosep) for peptide desalting and concentration, following the manufacturer’s instructions.

### Liquid-Chromatography tandem Mass Spectrometry (LC-MS/MS)

Data were acquired on a Fusion Lumos Tribrid Mass Spectrometer (Thermo Scientific) equipped with electrospray ionization source, coupled to a Evosep nano-LC system. Samples were run with a 15 spd (∼88 min) method with a flowrate of 220 nL/min in DIA mode. Buffer A consisted of H_2_O/0.1% FA and Buffer B was Acetonitrile/0.1% FA (LC-MS grade, Fisher Scientific). Peptides were separated by reversed-phase HPLC on an in-house packed analytical capillary column, with dimensions 25 cm, 150 nm internal diameter, with BEH 1.7 µm C18 resin (Waters). Peptides were eluted from the column tip and nanosprayed into the mass spectrometer, with spray voltage set to 2 kV at the back of the column. Collision energy (CE) was fixed at 30%, and peptides were fragmented with High-Energy Collisional Dissociation (HCD) in the orbitrap. Data were acquired with a 15 spd LC method (∼88-minute gradient) with a flowrate of 220 nL/min in DIA mode.

The DIA method consisted of 60 fixed-width, sequential isolation windows of width 10 *m*/*z* ranging from 300 – 900 *m*/*z* and performed in the quadrupole. Default charge state was +1, MS data type was set to profile and AGC target was set to 4e^5^ ions. Mass offset was set to 1.0005 with a 1 *m*/*z* overlap between windows. MS and MS/MS scans were performed in the orbitrap, with a resolution of 120K and 15K, respectively. MS/MS scans were performed over a range of 200 – 2000 *m*/*z*. Maximum injection time was 100 ms for MS and MS/MS scans. The MS/MS AGC target was set to 1e^6^ ions and data type was centroid.

### Data analysis

Raw MS data were converted to ‘.dia’ format and searched in DIA-NN v 2.1.0 ^72^ at 1% FDR against an *in silico* predicted library generated using the *Mus musculus* reference proteome (downloaded from Uniprot on 3 March 2025; 21,803 sequences). The settings were as follows: Protease -Trypsin/P; Missed Cleavages – 1; Max number of mods – 3; N-term excision, C carbamidomethylation, Ox(M) were enabled; MBR was disabled; --export-quant was enabled; all other settings were as default. The report file was loaded into R and the data were additionally filtered at Q.Value, PG.Q.Value, and Global.Q.Value ≤ 0.01. The diann_maxlfq function from the R package ‘diann’ was used to obtain aggregated protein quantities from peptide information. For all comparisons, proteins were considered detected if quantified in ≥ 2 replicates within each group, unless otherwise stated. The R package ‘enrichR’ (v 3.4) was used for enrichment analysis ^73^, and the online web tool for STRING protein network analysis (https://string-db.org/) was used to perform protein-protein interaction network analysis and generate all associated figures/plots. Post-hoc power analysis revealed >0.99 power for large effect sizes but only 0.38 power for moderate effects, indicating our protein lists represent minimum estimates of true group differences.

For identification of the IGG2B protein corresponding to the IL-6R antibody, we performed a separate DIA-NN search against the rat proteome (*Rattus norvegicus*; downloaded from Uniprot on 12 September 2024: 21,800 entries) with the same search settings. IGG2B peptide sequences were extracted from the rat search results, and three unique peptide sequences were identified (Supplementary file 2). The DILLISQNAK peptide was identified as unique to rat (i.e., non-homologous between mouse and rat), while the other two peptides were homologous between species. To visualize this peptide, it was copied into Skyline ^74^ and an *in silico* spectral library was generated using within Skyline (v 25.1.0.237, MacCoss Lab, Department of Genome Sciences, University of Washington, WA) using koina (intensity model: Prosit_2020_intensity_HCD; iRT model: AlphaPept_rt_generic). All settings were set as default for centroid/orbitrap data, and the DIA window information was imported using one of the results files. All the results files were then imported using a retention time (RT) tolerance of 10 min, and the spectra compared to the DIA-NN output for *m*/*z* and RT matching. The peak areas and associated information were extracted as confirmation of peptide identity (see Supplementary files 3 and 4).

## Supporting Information

All R scripts used for data analysis and figure generation can be found at the GitHub repository https://github.com/NataliePTurner/MouseCSF (*private until published*). All MS data raw files and sample annotations list have been uploaded to the MassIVE repository (https://massive.ucsd.edu/ProteoSAFe/static/massive.jsp), with the dataset identifier C5GX4573B. The final filtered quantitative MS data file containing protein IDs, *q* values and quantities can be found as Supplementary File 1. Supplementary file 2 contains the list of IGG2B peptide sequences identified using the rat proteome and filtered at 1% FDR. Supplementary file 3 contains the exported results from Skyline for the DILLISQNAK peptide, and Supplementary file 4 contains histograms of the DILLISQNAK peak areas normalized to total area for each sample.

## Acknowledgements

Support for this study was provided by National Institute on Alcohol Abuse and Alcoholism grants: T32 AA007456, AA013498 (MR, AJR, MB), P60 AA006420 (MR, JY, AJR), AA017447 (MR), AA029841 (MR), AA021491 (MR) and the Schimmel Family Chair. The authors thank Pauravi Gandhi for technical assistance with sample collection.

## Author contributions: CRediT

**Natalie P Turner**: Conceptualization, Methodology, Formal Analysis, Investigation, Data Curation, Writing – Original Draft, Visualization. **Michal Bajo**: Methodology, Investigation, Writing – Review & Editing, Visualization. **Amanda Roberts**: Methodology, Validation, Resources, Writing -Review & Editing. **Marisa Roberto**: Conceptualization, Methodology, Investigation, Writing – Review & Editing, Resources, Project administration, Funding acquisition. **John R Yates III**: Writing – Review & Editing, Resources, Project administration, Funding acquisition.

## Declaration of generative AI and AI-assisted technologies in the manuscript preparation process

During the preparation of this work the authors used Claude.ai Sonnet (v4.0) to write R code used to generate Figure 2, perform power analysis, and summarize the enrichment analysis data. After using this tool, the authors reviewed and edited the content as needed and take full responsibility for the content of the published article.

