## Supplementary file 4 for "The effects of Alcohol Dependence on the CSF Proteome in Mice: Evidence for Blood-Brain Barrier Dysfunction and Neuroinflammation"

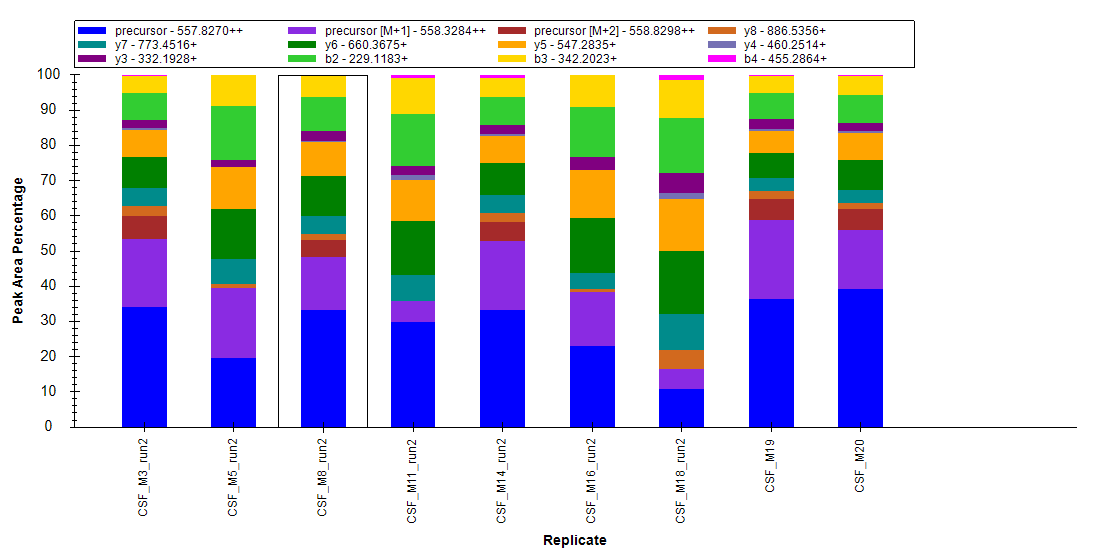

**Figure S1:** Peak area percentages of the IGG2B_RAT peptide DILLISQNAK (normalized to total peak area). Legend indicates ion types.
